# The Wisconsin Registry for Alzheimer’s Prevention: A Review of findings and current directions

**DOI:** 10.1101/206839

**Authors:** Sterling C. Johnson, Rebecca L. Koscik, Erin M. Jonaitis, Lindsay R. Clark, Kimberly D. Mueller, Sara E. Berman, Barbara B. Bendlin, Corinne D. Engelman, Ozioma C. Okonkwo, Kirk J. Hogan, Sanjay Asthana, Cynthia M. Carlsson, Bruce P. Hermann, Mark A. Sager

## Abstract

The Wisconsin Registry for Alzheimer’s Prevention (WRAP) is a longitudinal observational cohort study enriched with persons with a parental history (PH) of probable Alzheimer’s Disease (AD) dementia. Since late 2001, WRAP has enrolled 1,561 people at a mean baseline age of 54. Participants return for a second visit four years after baseline and subsequent visits occur every two years. Eighty-one percent (1270) of participants remain active in the study at a current mean age of 64 and 9 years of follow-up. Serially assessed cognition, self-reported medical and lifestyle histories (*e.g*. diet, physical and cognitive activity, sleep, and mood), laboratory tests, genetics, and linked studies comprising molecular imaging, structural imaging and cerebrospinal fluid data, have yielded many important findings. In this cohort, PH of probable AD is associated with 46% *APOE* ε4 positivity, more than twice the rate of 22% among persons without PH. Subclinical or worse cognitive decline relative to internal normative data has been observed in 17.6% of the cohort. Twenty-eight percent exhibit amyloid and/or tau positivity. Biomarker elevations, but not *APOE* or PH status, are associated with cognitive decline. Salutary health and lifestyle factors are associated with better cognition and brain structure, and lower AD pathophysiologic burden. Of paramount importance is establishing the amyloid and tau AD endophenotypes to which cognitive outcomes can be linked. Such data will provide new knowledge on the early temporal course of AD pathophysiology and inform the design of secondary prevention clinical trials.

## INTRODUCTION

While it is widely recognized that Alzheimer’s Disease (AD) has an extended preclinical stage, the cognitive and neuropathobiological course of changes in late-middle-aged people who may later develop AD dementia are relatively unknown [1]. Such knowledge is crucial if AD is to be identified in its inchoate form, its pathogenesis illuminated, and the tempo and predictors of its progression characterized as a predicate to successful prevention trials.

The Wisconsin Registry for Alzheimer’s Prevention (WRAP), established in 2001 [2], is a longitudinal observational cohort of participants who enrolled at mid-life (mean age 54), and that is enriched with risk for late onset AD due to parental history (PH) of AD dementia. The cohort also serves as a registry for linked studies. The overarching goals of the study shown in Table 1 are to identify early cognitive decline, and to characterize mid-life factors associated with such decline and the contributing underlying biomarkers of AD and related pathology. The present contribution updates the initial description of the cohort, study design and protocol [2], and provides new data on the effects of family history, APOE genotype and AD biomarkers on longitudinal cognitive decline over time. Key study findings are summarized and future directions are presented.

**Table 1.**
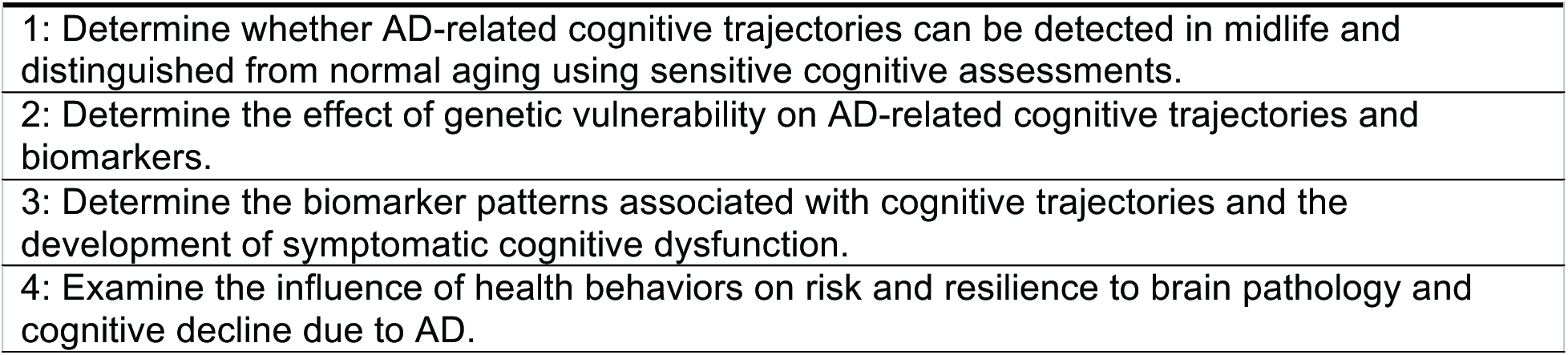
The Major Goals of the WRAP study

## METHODS

### Study Design

To the present, 1,561 participants have enrolled over a continuing enrollment window. Recruitment sources included memory clinics in which a parent was diagnosed or treated, limited radio and newspaper advertisements and word of mouth. Participants generally meet the following inclusion criteria at study entry: age 40 to 65; fluent English speaker; visual and auditory acuity adequate for neuropsychological testing; good health with no diseases expected to interfere with study participation over time. Participants are excluded from enrollment if they have a prior diagnosis of dementia or evidence of dementia at baseline testing (1 was excluded due to baseline dementia). The baseline mean age is 54, 73% have a parent with AD dementia, and 40% of the total sample are *APOE* ε4 carriers (46% of the PH+ participants and 22% of the PHparticipants).

#### Determination of parental history of AD

The characteristic of parental history of AD (PH+) is defined as having at least one biological parent diagnosed with dementia due to probable AD based on the NINDS-ADRDA criteria [3]. Three general methods were used to determine PH. First, direct diagnosis of the parent from study physicians or affiliated faculty, or where medical records for the affected parent were available, a panel of study investigators reviewed the parent’s clinical evaluation for dementia to determine whether evidence was sufficient to diagnose probable AD. Second, neuropathological confirmation of AD in the affected parent. Third, in the absence of sufficient prior information, a Dementia Questionnaire [DQ; 4] was conducted with the adult child regarding the parent’s dementia history and course. The DQ asks about the type of dementia symptoms, the course of progression and the presence or absence of co-morbid conditions that could explain or contribute to the symptoms. Diagnostic classifications based on the DQ show very high sensitivity (100%) and specificity (90%) compared to clinical diagnosis [5]. Eight percent of PH subjects initially qualified for study entry by a parental autopsy; 83% by medical record review or expert physician diagnosis; and 9% by DQ. Less than 1% were self-report.

#### A comparison group without PH of AD

To understand the role of PH, recruitment of additional participants without PH of probable AD dementia began in 2004. This group now consists of 421 persons who by self-report did not have a parent with dementia due to AD or related cause, and who in general have a mother who survived to at least age 75 and a father to at least age 70 without dementia.

Because parental status changes over time, it is reassessed at each visit and updated as necessary (*e.g.* in the case that a previously non-demented parent later developed dementia or, rarely, a parent whose dementia was presumed due to AD was later found by autopsy to be another pathology).

### Study Visit Procedures

Participants are followed at regular intervals with detailed in-person assessments, questionnaires, and blood collection occurring at each study visit. The first follow up is approximately 4 years after baseline, and further follow up visits are approximately every two years. Persons will remain in the study until age 85, unless they withdraw, convert to dementia, or develop another illness precluding participation or accurate assessment of cognition. Each visit requires approximately five hours and comprises the assessments shown in Table 2 *i.e.*, cognitive measurement, anthropometric measures, laboratory tests, and questionnaire ratings completed by the participant and an informant including the Quick Dementia Rating System or Clinical Dementia Rating [6]. Reliability and consistency of cognitive testing is established through regular review of aspects of testing procedures at team meetings, biannual individual observations of test administration, through adherence to a standardized manual of procedures, and through blinded rescoring by a separate rater (20% annually for each psychometrist).

#### Consent for Brain donation

Neuropathologic confirmation is critical for linking cognitive trajectories to disease-related endpoints. Accordingly, participants are encouraged to enroll in the Wisconsin Brain Donation Program which is administered by the Neuropathology Core of the Wisconsin Alzheimer’s Disease Research Center. Brain bank enrollment has not been an entry criterion. However, since 2015 brain donation has been systematically discussed with participants at each visit, and educational material on the value of brain donation is regularly offered at WRAP’s statewide series of information sessions and in semi-annual newsletters.

#### Identifying subtle, preclinical impairment

A critical issue for the field is development and validation of optimal methodology for identifying early cognitive decline and impairment [7]. Simple single-test thresholds are insufficient [8] and available published norms used to define ‘impaired’ and ‘normal’ performances on neuropsychological tests in persons age ̴55+ may be confounded by unintended inclusion of individuals with incipient disease in the normative samples of those tests [9-11], thereby reducing sensitivity to subtle dysfunction [10]. Moreover, thresholds and norms may have been validated by others in populations of uncertain relevance to the cohort under investigation. To avoid these potential confounders, and to enhance sensitivity to preclinical decline, we developed a ‘robust’ norms approach in which internal normative distributions for cognitive factor scores [12] and individual test scores [13], are generated, where ‘robust’ indicates that the normative group is non-declining over time. In Koscik *et al*. [12], deficits on multiple visits via algorithmic criteria were required as evidence of “psychometric MCI” while in Clark *et al*. [13], deficits on multiple tests within a specific domain were required to identify persons with psychometric MCI. In practice, and to ensure that these approaches are not falsely over-identifying people with abnormal cognition, we incorporate these algorithms into our consensus review process as described in the next section.

#### Classification of cognitive status

If cognitive abnormalities are detected by algorithm on neuropsychological tests, data from participant visits are brought to a consensus review committee consisting of dementia specialist physicians, neuropsychologists and nurse practitioners for in-depth review. Thresholds for committee review include performance greater than 1.5 SD below robust internal norms adjusting for age, gender, and literacy-level [12, 13], self or informant report of cognitive or functional decline on the Clinical Dementia Rating (CDR), the Quick Dementia Rating Scale (QDRS), the Informant Questionnaire on Cognitive Decline in the Elderly (IQCODE), or instrumental activities of daily living; or threshold-specific absolute scores on key tests (*e.g.* Wechsler Memory Scale-Revised Logical Memory-II ≤17; Rey Auditory Verbal Learning Test delayed recall ≤5; or Mini Mental State Exam ≤26). The consensus committee assesses cognitive performance at all prior visits in order to detect intra-individual changes over time, and analyzes pertinent findings from the neurological and physical exam, medical and social history, and self- and informant-surveys of mood, cognition and functional status. The diagnosis of “mild cognitive impairment (MCI)” is based on National Institute on Aging–Alzheimer’s Association criteria [7] and requires a) patient or informant concern regarding change in cognition, b) unambiguous impairment in one or more cognitive domains, c) not meeting criteria for dementia. The experimental category of “early MCI” is assigned if there is lower than expected objective performance (typically >1.5 SD below internal robust norms), but few or no subjective cognitive complaints or clinically significant deficits [for further discussion see 14]. This category broadly corresponds to clinical stage 2 in the 2018 diagnostic framework [15].

**Table 2.**
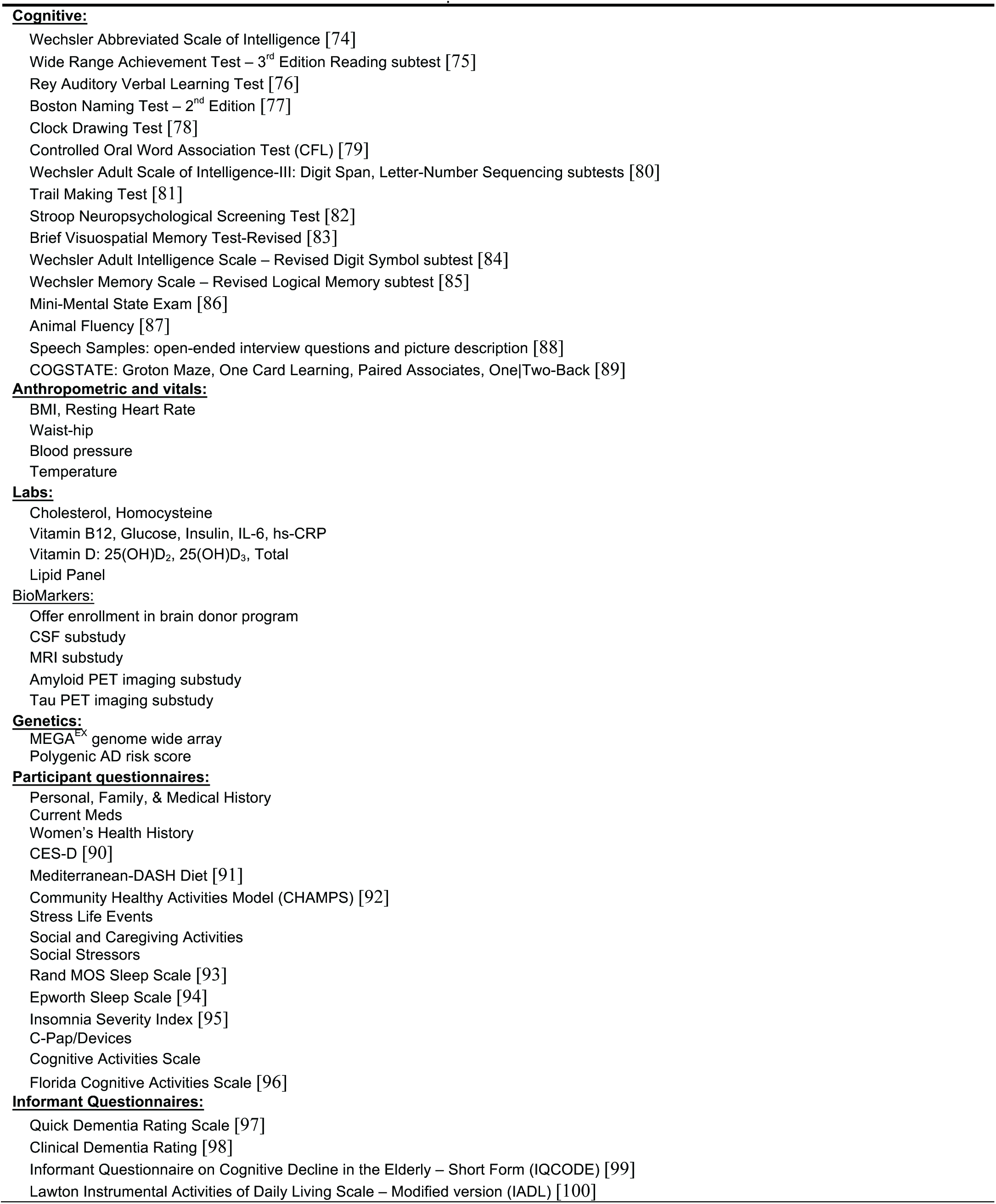
List of Procedures and tests in the current protocol

#### Biomarker and genetics procedures

A diversity of MRI, molecular PET and CSF biomarkers have been acquired from subsets of participants as funding permits (see **Supplemental Table** for sample sizes of each to date). Serial MRI’s and LP’s will be obtained from approximately 60% of WRAP participants over the next 5 years and serial amyloid and tau imaging will be obtained from approximately 30% with current and projected funding.

#### MRI protocol

In 2009 a standardized MRI protocol was implemented across the Wisconsin Alzheimer’s Disease Research Center (ADRC) to include all WRAP-linked studies. The standard protocol comprises an inversion recovery prepared T1-weighted 3D volume structural scan, a T2-FLAIR 3D volume to assess white matter hyperintensities, pseudo-continuous arterial spin labelling of cerebral blood flow, multishell diffusion weighted imaging to assess white matter integrity and structural connectivity, and 4D-flow imaging to assess intracranial blood flow and vessel stiffness.

#### CSF Collection and Analyses

Cerebrospinal fluid samples are collected in the core WRAP study as well as in linked studies. A center-wide standard pre-analytic protocol is 10 used to collect approximately 22ml of CSF which are subsequently gently mixed to remove collection gradients, partitioned into .5ml aliquots in 1.0ml polypropylene tubes, and stored at −80°C. Assayed analytes include total tau, hyper-phosphorelated tau (Ptau181), beta amyloid1-42 (Aβ42), beta amyloid1-40 (Aβ40), YKL40, neurofilament light-chain protein (NFL) and neurogranin [see, for example, 16, 17-23].

#### Molecular Amyloid and Tau Imaging

Amyloid imaging is conducted with [C-11] Pittsburgh Compound-B (PiB) positron emission tomography (PET) using a dynamic 70-minute protocol. For a full description of the PiB protocol, refer to [24]. All participants who undergo amyloid imaging are invited to undergo tau PET imaging with [F-18]MK6240 from ̴70-110 min post-injection. Derived maps of AD pathology burden are analyzed for longitudinal change.

#### Genetics

*APOE* ε2/ε3/ε4, 20 common genetic variants from the International Genomics of Alzheimer’s Project consortium [25], and low frequency variants in *TREM2* [26, 27] and *PLD3* [28] were genotyped using competitive allele-specific PCR based KASP genotyping assays (LGC Genomics, Beverly, MA). Duplicate quality control (QC) samples had 99.9% concordance. Cross-validation of *APOE* genotypes with prior assays was 99.7% concordant. Various polygenic risk scores are derived in which the contribution of each SNP to the score is weighted by its risk odds ratio [29].

More recently genome-wide genotyping was performed using the Illumina Infinium Expanded Multi-Ethnic Genotyping Array (MEGA^EX^) containing approximately 1.7 million genetic markers.

## REVIEW OF SELECT STUDY FINDINGS AND NEW RESULTS

Accrual: Figure 1 depicts participant accrual from November 2001 through May 2017. Accrual is shown by visit number together with mean age at each visit. The rolling recruitment window means that individual participants have a different number of followup visits to date depending on how long they have been in the study. Sixth visits began in late 2016. Over 5600 study visits have occurred since inception, and over 3000 visits are projected over the next five years. Retention is 82% over the 16-year study period.

Descriptive information: Baseline characteristics of the WRAP sample, including demographics, medical history, and cognition, are described in Table 3. In keeping with the original study design, Table 3 is stratified by parental family history and *APOE* ε4 carrier status. Histograms showing the sample ages at baseline and at last visit are provided in Figure 2.

**Figure 1.**
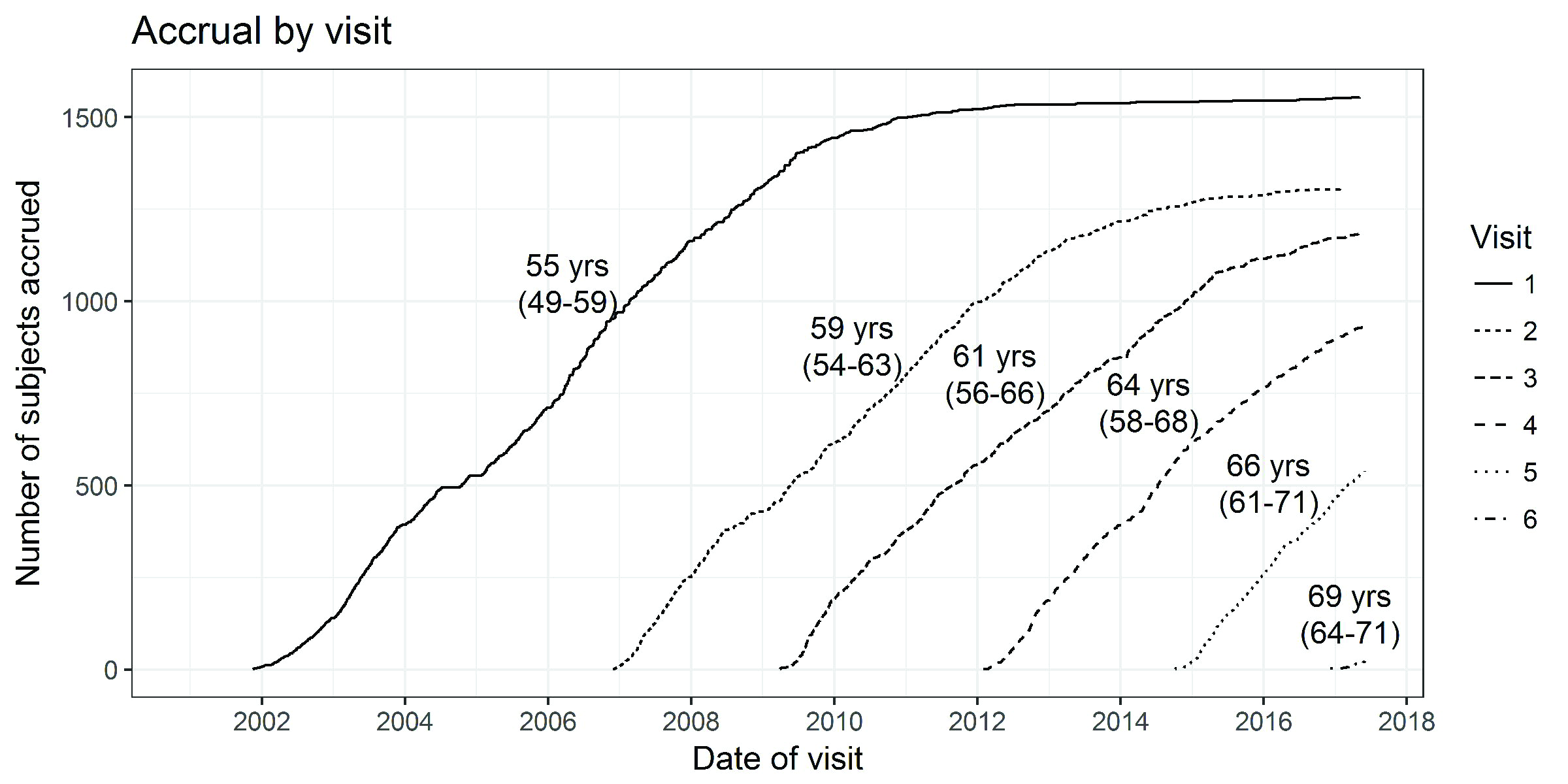
Accrual by study visit. Lines representing each visit are annotated with the median participant age (interquartile range) at that visit.

**Figure 2.**
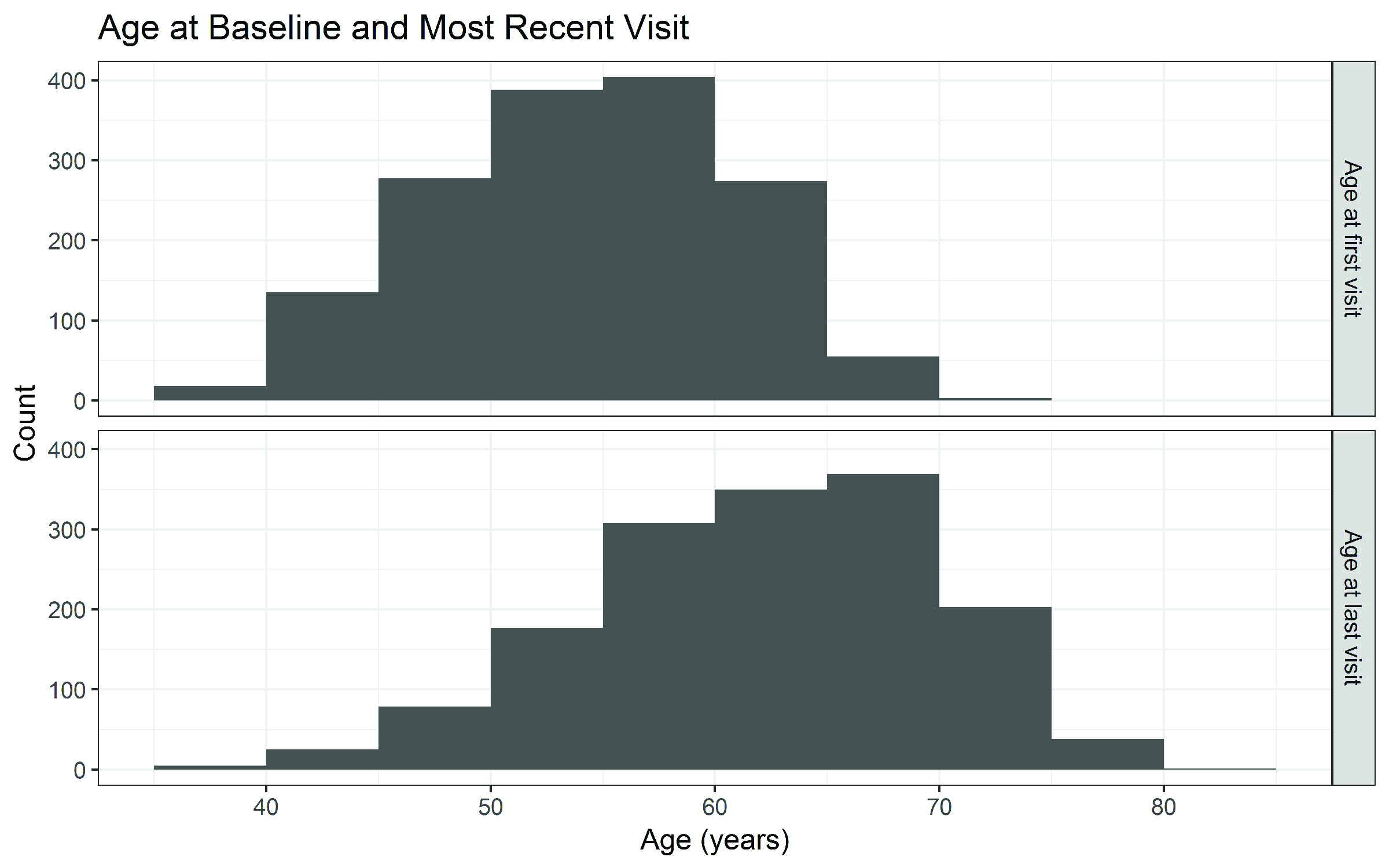
Histograms displaying the WRAP sample age distribution at baseline (upper panel) and most recent visit (lower panel). In the data for the lower panel, 15.9% of the sample have completed only one visit, so the reported numbers reflect age at Visit 1; for 7.78%, age at Visit 2; 16.4% for Visit 3; 25.2% for Visit 4; 33.2% for Visit 5; and 1.41% for Visit 6, respectively.

**Table 3.**
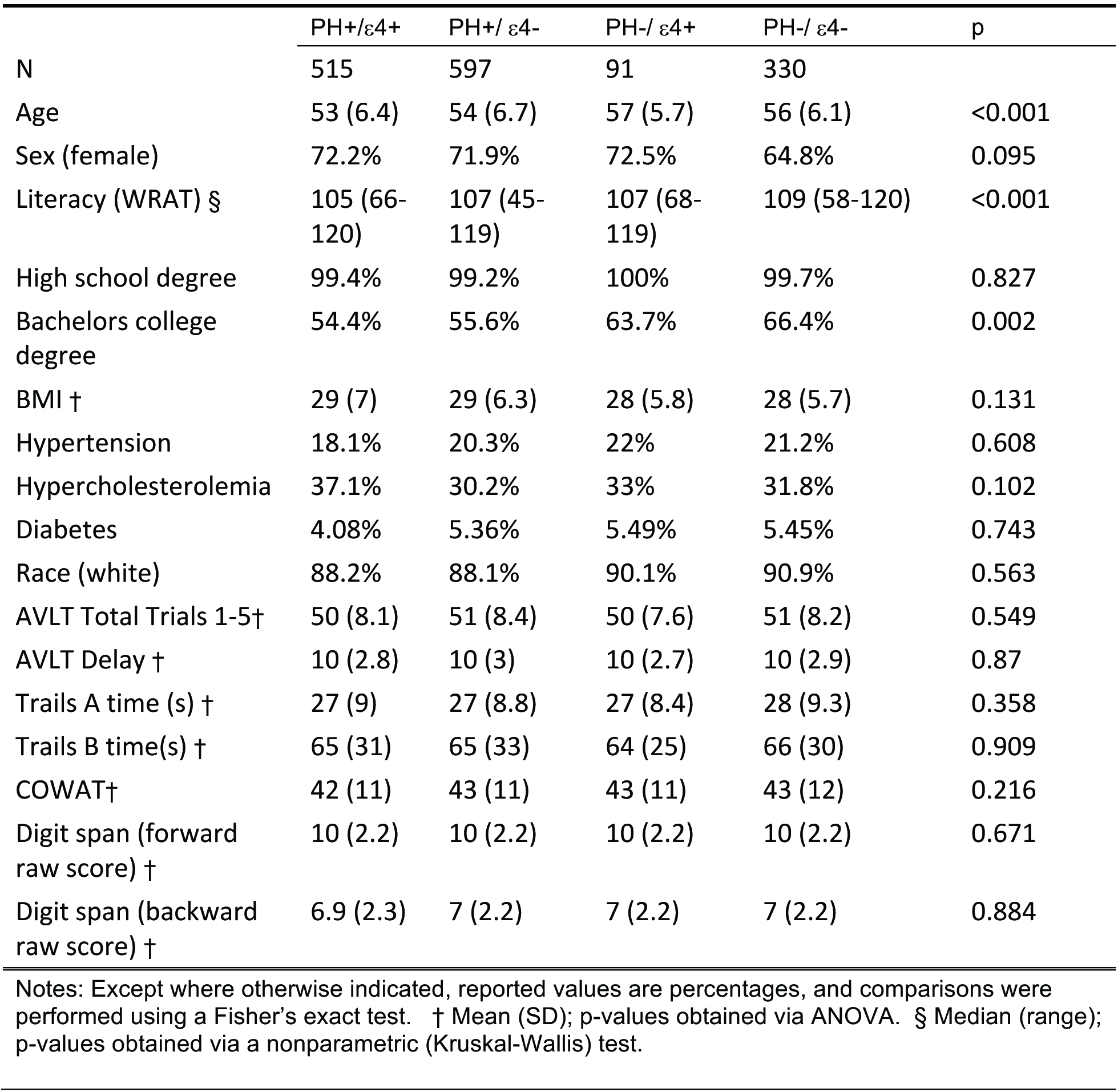
Baseline demographic and health characteristics grouped by parental history and *APOE* ε4 status.

### Results and discussion related to Goal 1

Determine whether AD-dementia related cognitive trajectories can be detected in midlife and distinguished from normal aging using sensitive cognitive assessments.

#### Previously published findings

As noted in the Methods section, we developed and evaluated two cross-sectional algorithms for identifying performance that is below robust internal norms using factor scores [12, 30] and using individual test scores [13]. Compared to the use of published norms in the same cohort, both psychometric approaches demonstrate improved sensitivity to small but consequential decrements of cognitive function. With the factor score approach [12], we observed greater longitudinal cognitive decline among the 13% of the cohort who were classified as psychometric MCI compared do those who were cognitively normal. Using this approach, we also showed that psychometric MCI is associated with greater dysfunction in connected language [31] and verbal fluency [32]. Using the cognitive test approach [13], the false positive rate was reduced by requiring a pattern of lower than expected performance on multiple tests within or across domains to be affected in order for the participant to be designated as having MCI [similar to 33, 34]. With use of this “multi-test, single visit” approach, 18% of participants were classified as having psychometric MCI. These algorithmic approaches may serve in their own right as intermediate outcomes, and are used to inform research visit-by-visit diagnoses made by the diagnostic consensus committee. In analyses of consensus committee classified cognitive status, higher intra-individual cognitive variability at baseline (IICV) predicted impaired cognitive status 8-10 years later [14].

##### New results

15.2% of the cohort met early MCI criteria at their last study visit while 2.3% met criteria for MCI and 0.1% met criteria for dementia due to AD. Prevalence for early MCI at the most recent visit was associated with age, ranging from approximately 6 percent for the youngest participants to 20 percent for those over 70. Prevalence of early MCI, MCI, and dementia by last visit age are shown in Figure 3.

In order to determine whether demographic and risk features differ by the clinical status and age groups shown Figure 3, we conducted Cochran-Mantel-Haenszel (CMH) tests for independence for the variables *APOE* ε4 status, PH of AD status, sex and presence of a college degree on the outcome categorical variable of cognitive status (cognitively normal, early MCI, and MCI/dementia (combined because there are so few dementia cases), after adjusting for age category. No significant associations between cognitive status and APOE, PH of AD or education level (<BA vs >= BA) were evident after adjusting for age grouping. Sex and cognitive status were not independent (CMH general association test *p*=.0018) such that men (which comprised 28.6% of the whole sample) were more likely to have eMCI (n=90 men; 38.3%) or MCI/Dementia (n=16 men; 43.2%). The cohort is relatively young and these associations may change over time.

The designations of ‘early MCI’ and ‘psychometric MCI’ have been used here and in other recent papers to describe cognitive decline that is not sufficiently severe to warrant a diagnosis of MCI. This intermediate stage may have predictive value for identifying persons at risk of progression to MCI or dementia. Specifically, of those who were early MCI at visit 1, 11.4% progressed to a clinical status at their last study visit compared with 2.0% of those who were cognitively normal (*χ*^2^(1)= 35.4, p<.0001). Whether persons with emerging impairment also have greater biomarker signs of AD or vascular pathology is a topic of ongoing investigation. In the parlance of the 2018 research diagnostic framework these intermediate categories largely overlap with clinical stage 2 which connotes cognitively unimpaired with decline from a prior baseline [15]. Future work will implement the new research criteria in WRAP’s diagnostic processes.

**Figure 3.**
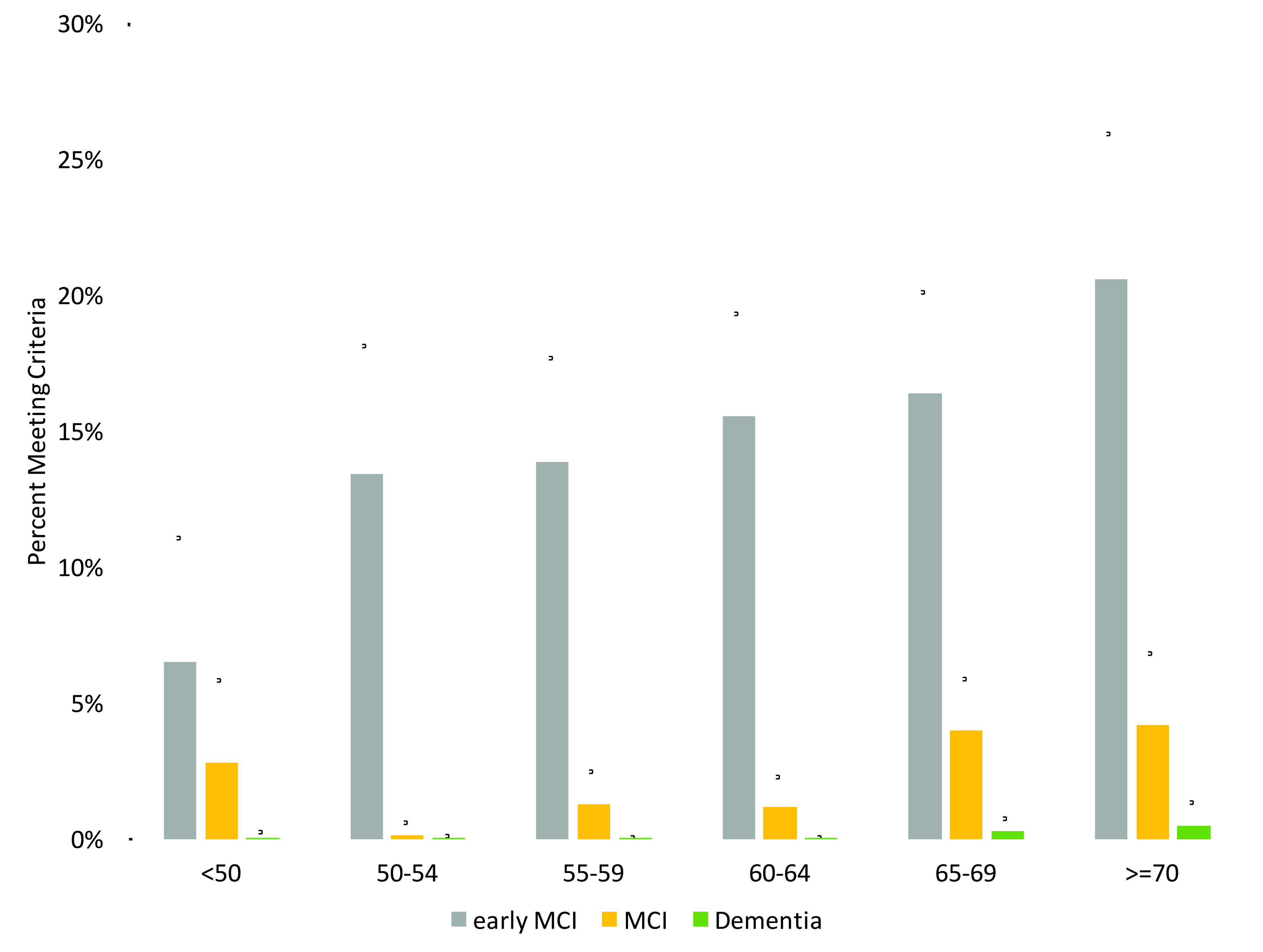
Cognitive statuses at last study visit, by 5-year age increments. The denominators for each age range were 108 (<50), 194 (50-54), 310 (55-59), 341(60-64), 378 (65-69), 214(>=70).

### Results and discussion related to Goal 2

Characterize the effect of parental history and genetic vulnerability on AD-related cognitive trajectories.

#### Previously published findings

In addition to identifying specific individuals with cognitive impairment, the study seeks to understand how PH of AD and *APOE* ε4 status (both major risk factors) affect cognitive trajectories from midlife. Earlier cross-sectional analyses of WRAP cognitive data suggested modest relationships between cognitive performance and genetic and/or parental history risk factors. For example, at the first visit, PH+ and PH- participants had similarly high scores on a list-learning task, but PH+ participants relied more heavily on recent list items, suggesting greater difficulty with consolidation [35].

As expected, WRAP PH+ participants are more than twice as likely to be *APOE* ε4 positive (see Table 2). They are also more likely to carry the *TREM2* T risk allele [36]. As well, genetic risk features for AD may interact with one another. Two variant alleles of the *ABCA7* gene are associated with worse memory and executive function scores in participants with no *APOE*-ε4 alleles, but with better scores in those with 1 or 2 *APOE*ε4 alleles [37]. To the present, longitudinal comparisons have not detected effects of these genetic markers on cognitive trajectories [36, 37]. Boots *et al*. [38] examined the influence of the brain-derived neurotrophic factor (*BDNF*) Val66Met polymorphism on cognitive trajectories and PET amyloid load. Compared with Val carriers, *BDNF* Met carriers exhibit greater decline over time on the Verbal Learning and Memory factor score. In the subset with amyloid imaging, amyloid burden modified the relationship such that those with high amyloid burden who were also *BDNF* Met carriers exhibited steepest cognitive decline. The aggregate effects of 21 genetic risk alleles on cognition *via* a polygenic risk score (PRS) identified modest negative effects on working memory performance, but not other domains [29]. Darst and colleagues [29] also examined PRSs specific to causal pathways implicated in AD. Gene clusters affecting Aβ clearance and cholesterol metabolism were strongly predictive of CSF Aβ_42_, CSF Aβ_42_/Aβ_40_, and PiB amyloid burden, as was *APOE* when considered as an independent predictor on its own [see also 23].

##### New results

To explore the effects of PH and *APOE* on longitudinal cognitive trajectories, we modeled the trajectories with a linear mixed effects model [39, 40] using random intercepts at the family and participant levels, and a random age slope at the participant level. Fifty-nine participants reporting neurological diagnoses at baseline were excluded from these analyses. Baseline characteristics of this subset were virtually identical to those in Table 3. Covariates included age (linear and quadratic age terms were tested), sex, race, education level, and baseline literacy (WRAT-III Reading), and the number of prior exposures to the neuropsychological test battery [41]. The effects of interest are: (1) the main effects of *APOE* and PH (baseline effects); (2) their interaction with age (longitudinal effects); and (3) their interaction with prior test exposure, that accounts for the differential benefit from practice. Table 4 lists estimates for each of these terms for Verbal Learning and Memory, an important composite outcome for early cognitive change [30]. Predicted trajectories with age are plotted by *APOE* and PH status in Figure 4.

**Figure 4.**
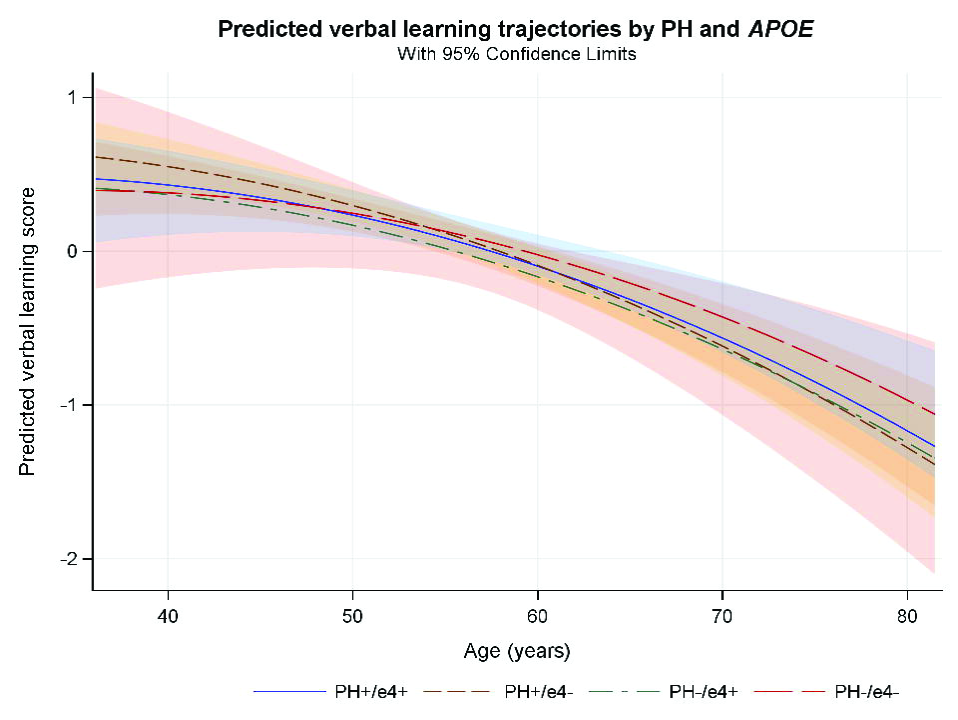
Verbal memory decline by parental history of dementia due to probable AD and *APOE* ε4 binary status (presence/absence of PH and of *APOE* ε4). All four groups declined with age, but no significant differences between groups are evident.

**Table 4.**
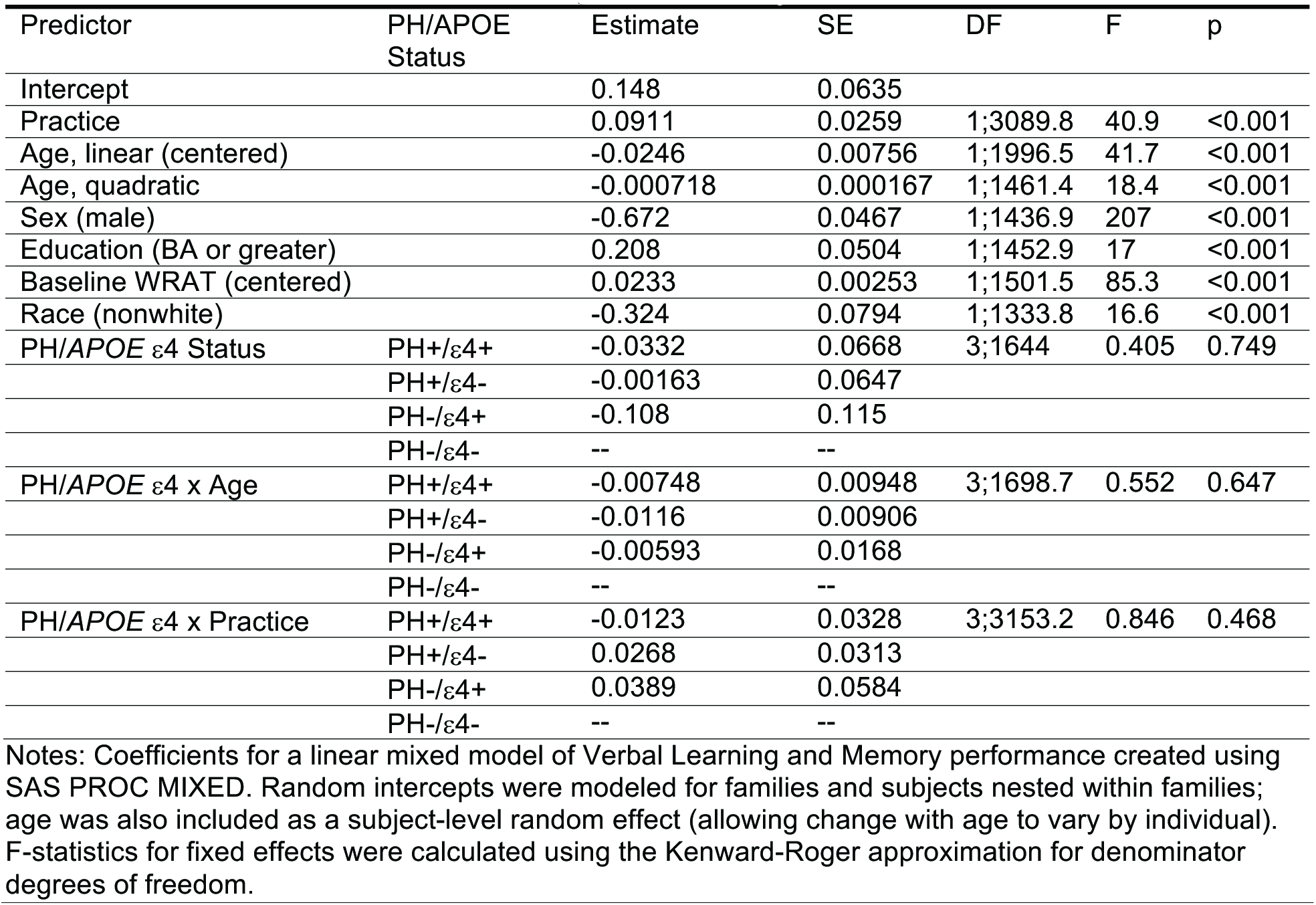
Results of Mixed effects model of cognitive decline by *APOE* ε4 and PH status.

Although small decrements in performance occur with increasing age, little evidence is observed in these data for effects of *APOE* or PH on cognitive performance, either in the main analysis or in secondary analyses that (a) excluded participants whose parents developed AD after age 75 and (b) modeled cognition as a function of *APOE* genetic risk score rather than an *APOE* ε4 binary status variable [29]. The lack of *APOE* or PH risk effects on cognition is unlikely to be due to lack of power. Prospective power calculations using Monte Carlo simulations (k=1000) indicated power of 0.95 or greater to detect small main effects (mean performance approx. 0.2 SD lower for the highest-risk group), and age interactions (age-related slope approx. 20% steeper for the highest-risk group). Because the cohort is still relatively young (current mean age 64) and will continue to be followed to determine if an effect unfolds with older age, we caution that the current reported absence of an effect in these data should not be accepted as definitive for the WRAP cohort.

For the subsample who underwent amyloid imaging or CSF collection and on whom Aβ42 assay results were available (N=211), we classified each participant as amyloid-positive (n=62; 29%) or amyloid-negative (n=149; 71%) using cutoffs described elsewhere [17, 42]. In this subset, amyloid positivity was significantly related to PH (Table 5; X^2^ = 14.22, p = 0.003). However, follow-up tests indicated the association was largely explained by *APOE* ε4 carrier status (X^2^ = 12.82, p < 0.001) rather than an independent effect of PH specifically (X^2^ = 0.88, p = 0.35). The significant relationship between *APOE* ε4 status and amyloid status holds even after controlling for age at the date of biomarker assessment (logistic regression: β_*APOE*_ = 1.263, p < 0.001). In a smaller subsample for whom longitudinal amyloid (PiB) data were available (N=142), conversion from amyloid-negative to amyloid-positive was associated with carriage of at least one *APOE* ε4 allele (X^2^ = 4.24, p = 0.04) but not with PH, baseline age, gender, or consensus conference diagnosis (all p > 0.10). Together with the null cognitive findings, these results are consistent with a prevailing biomarker model in which AD pathophysiology precedes cognitive change [1, 43].

**Table 5.**
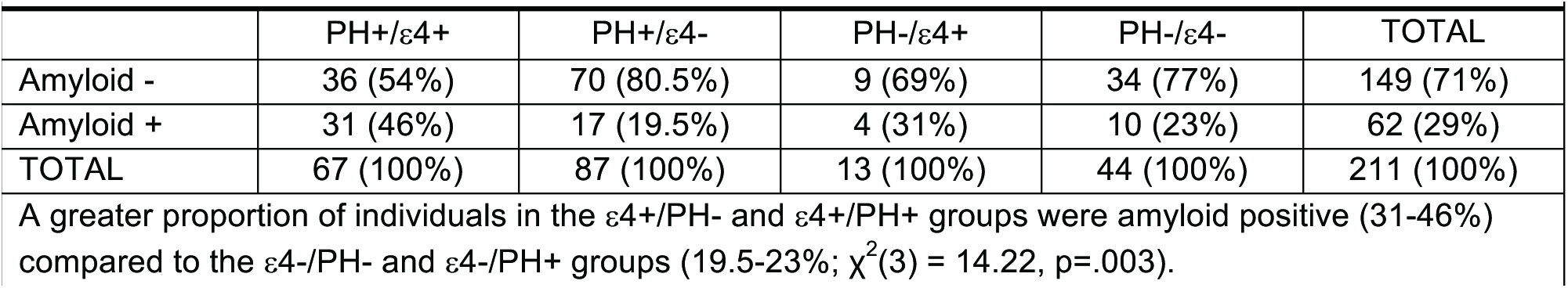
Risk (PH and *APOE* status) versus amyloid positivity.

### Results and Discussion related to Goal 3

Determine the biomarker profiles associated with cognitive decline and the development of symptomatic cognitive dysfunction.

The WRAP study spans a period of scientific development in which risk factors like APOE status and PH have given way to more direct biomarkers of AD pathology for characterizing the preclinical stages of AD [1, 43-46], and less specific brain markers of function and health such as functional and structural MRI have become gauges of the effect of AD pathophysiology on brain function and neurodegeneration. Early structural and functional MRI studies in WRAP (uninformed at the time by AD pathology biomarkers of amyloid and tau) found differences in cerebral activity during memory tasks [47-49] diffusion tensor imaging [50] and hippocampal morphology [51] as a function of PH or APOE ε4 status. These were interpreted as potentially related to differential AD pathophysiological processes in the PH group, though there was nothing specific to AD-pathology about the imaging at the time or about the risk features by which the participants were stratified.

More recently, the effects of CSF Aβ42 and tau, and amyloid PET imaging using [C-11]PiB have been examined as indices of the presence or absence of AD pathology. In n=201 WRAP participants with [C-11]PiB amyloid PET data at a mean age of 61, there was no significant relationship with concurrent cognition [24]. This finding was not surprising as amyloid load is expected to increase prior to cognitive loss. Subsequently Clark et al. [52] tested cognitive changes over time with mixed effects models, and found that greater overall amyloid burden via PiB PET was associated with a greater decline on composite test performance in episodic memory and executive function. Similar results have been found with CSF derived estimates of amyloid and tau burden [42] suggesting that cognitive consequences of AD pathology may already be present reflecting the gradual accumulation of disease. Moreover, other data suggest that AD pathology in the late-midlife preclinical time frame may co-exist with atrophy and/or vascular and other diseases that have secondary effects on neural tissue. Racine *et al.* [17] used hierarchical clustering analysis of amyloid burden, tau burden, white matter hyper-intensities and hippocampal atrophy to categorize WRAP and comparable Wisconsin ADRC participants with imaging and CSF data. Four clusters emerged including: 1) participants with preclinical AD pathology who were predominantly positive for tau and amyloid; 2) participants with mixed vascular and AD pathology who exhibited white matter hyper-intensities as well as variable AD pathology; 3) participants with suspected non-AD pathology who exhibited atrophy but not Ab or tau pathology; and 4) participants with healthy aging who exhibited normal imaging and CSF biomarkers. The greatest decline on memory tests over time was observed in the preclinical AD cluster. Taken together, these findings indicate that biomarkers of AD pathology are sensitive to cognitive decline in late middle-aged WRAP participants. As the cohort ages and individuals develop age-related diseases, biomarker profiles will likely become more heterogeneous. Thus, models examining change in biomarkers and cognition over time must include more precise markers of other pathologies as they become available.

### Results and Discussion related to Goal 4

Characterize the influence of health behaviors on risk and resilience to brain pathology and cognitive decline due to underlying AD.

#### Previous findings

The effects of modifiable risk factors on cross-sectional cognition have been the target of multiple WRAP investigations. Cognitive activity throughout the lifespan measured by education level [22], job complexity [53], and self-reported current participation in stimulating activities such as games [54, 55], are associated with better performance in several cognitive domains. Participants with greater numbers of stressful life events perform worse on measures of cognitive speed and flexibility, and conversely, in participants with greater social support performance is better [56]. Of note, the protective effect of social support is diminished by presence of *APOE* ε4 [57]. Sleep adequacy in WRAP participants is associated with amyloid burden assessed with amyloid imaging [58] and CSF Aβ42 and tau levels [19]. Cardiovascular and metabolic fitness also appear to have a protective effect on gray matter, cerebral blood flow and episodic memory performance. Insulin resistance in particular is linked with cerebral atrophy, amyloid burden, CSF biomarkers of AD pathology, and lower cerebral glucose uptake [59-64].

#### Physical Activity (PA) and brain health

PA and related variables including cardiorespiratory fitness (CRF) are well-studied protective factors for AD dementia, with a recent evidence review identifying PA as the modifiable factor with the highest impact on reducing the national prevalence of this disease [65, but see also 66]. Cross-sectional WRAP publications elucidate the relationship between PA, cognition, brain structure, and neuropathological markers of AD among normal adults. In n=315 WRAP participants, Boots *et al.* [67] found that engagement in PA was associated with preserved volume in diverse brain regions including the medial and lateral temporal lobe and medial parietal lobe, together with reduced white matter ischemic lesions and with fewer memory complaints. Dougherty *et al.* [68] assessed PA *via* accelerometer in 91 WRAP participants, and found that those meeting recommended PA levels had greater temporal lobe regional volumes including the hippocampus compared to those who did not meet recommended activity levels. In addition to effects on brain structure and cognition, PA moderated the effect of age [69] and genetic risk factors [23] on AD pathophysiological biomarkers in WRAP participants. Specifically, Okonkwo *et al.* [69] report that a history of PA was associated with an attenuation of age-related alterations in Ab burden, cerebral glucose metabolism, and hippocampal volume. Similarly, Schultz *et al.* [23] found that CRF attenuated the adverse influence of cholesterol metabolism polygenetic risk on CSF biomarkers. Of note, the beneficial effect of PA on the brain substrates of cognitive health may depend on level of exercise intensity [70].

In aggregate these associational studies in WRAP participants suggest that modifiable factors such as physical and cognitive activity, glucose and metabolic regulation, stress and sleep may be avenues for interventions that enhance brain health and reduce the likelihood and severity of AD pathology.

### Future Directions

WRAP and its linked studies are charting the preclinical time course of Alzheimer’s disease. Ongoing WRAP investigations assess lifestyle, genetic risk and resilience factors along with longitudinal cognitive and clinical assessments to establish whether AD biomarker trajectories and cognitive trajectories can be identified in midlife. The associational findings from WRAP which have been partially summarized here now require further study to determine whether improving these health behaviors can result in measureable effect on AD biomarkers and brain and cognition health in late-midlife.

Whereas WRAP’s organizing theme in 2001 was risk enrichment due to parental history of dementia due to probable AD, focus has broadened in the ensuing years to AD biomarkers. In the present phase of WRAP, CSF and PET biomarkers of AD pathology are sought from all willing participants, funding permitting, as visits at 2-year intervals continue. Biannual (and now annual) information sessions with participants to share back what we are learning serve to educate participants on the overarching importance of biomarker enrollment and brain donation, and are also an effective retention component. Although PH as an enrichment factor has been supplanted in part by the capability to directly measure AD pathology *in vivo*, the experience of having a parent with AD dementia motivates many of our participants to remain in WRAP and take part in linked studies at a high level of volunteerism. A caveat is that the participant characteristics are biased toward and most generalizable to persons who have a parent with dementia due to probable AD by design. The cohort is also biased in other ways. Because WRAP is a self-selected sample of convenience, the majority of the cohort are Caucasian (88%), women (71%), and highly educated (mean 16 yrs). Increasing the ethnic diversity of WRAP participants and assuring that WRAP’s findings are generalizable to African Americans in particular is a current priority.

Data from the core WRAP protocol and from a subset of WRAP’s linked studies are accessible to qualified researchers *via* an online request form and data use agreement which can be linked from the Global Alzheimer’s Association Interactive Network website (www.gaain.org). Recent examples of data sharing collaborations include a study that found consistency in patterns of cognitive aging progression scores across the Baltimore Longitudinal Study on Aging and WRAP [71], a meta-analysis of *APOE* and sex on dementia incidence [72], and a study involving predictive algorithms for MCI in a consortium of five preclinical AD or adult children cohorts [73].

The WRAP observational longitudinal cohort is AD risk-enriched, and has been followed with detailed measurements since midlife. This is a time frame that is less well studied than older ages, but is nevertheless a critical epoch, as this is when AD pathology likely begins and when its trajectory may be modifiable through pharmacologic and/or lifestyle approaches.

## ACKNOWLEDGMENTS

This research was supported by the National Institutes of Health awards (R01 AG027161, R01 AG021155, R01 AG054047, R01 AG037639, P50 AG033514, UL1 TR000427, K23 AG045957, F30AG054115, T32 GM007507, and T32 GM008692) and grants from the Alzheimer’s Association, Extendicare Foundation, and Northwestern Mutual Life Foundation to the University of Wisconsin–Madison; and by donor funds including the the Wisconsin Alzheimer’s Institute Lou Holland Fund and contributions from anonymous donors. Portions of this research were supported by resources at the Wisconsin Alzheimer’s Institute, the Wisconsin Alzheimer’s Disease Research Center and the Geriatric Research Education and Clinical Center of the William S. Middleton Memorial Veterans Hospital, Madison, WI. Any opinions, findings, and conclusions or recommendations expressed in this material are those of the authors(s) and do not necessarily reflect the views of the NIH or the Veterans Administration. We gratefully acknowledge the WRAP study team who have carefully acquired the longitudinal data, and the WRAP participants who make this research possible.

